# Single cell transcriptome atlas of immune cells in human small intestine and in celiac disease

**DOI:** 10.1101/721258

**Authors:** Nader Atlasy, Anna Bujko, Peter B Brazda, Eva Janssen-Megens, Espen S. Bækkevold, Jørgen Jahnsen, Frode L. Jahnsen, Hendrik G. Stunnenberg

## Abstract

Celiac disease (CeD) is an autoimmune disorder in which ingestion of dietary gluten triggers an immune reaction in the small intestine^1,2^. The CeD lesion is characterized by crypt hyperplasia, villous atrophy and chronic inflammation with accumulation of leukocytes both in the lamina propria (LP) and in the epithelium^3^, which eventually leads to destruction of the intestinal epithelium^1^ and subsequent digestive complications and higher risk of non-hodgkin lymphoma^4^. A lifetime gluten-free diet is currently the only available treatment^5^. Gluten-specific LP CD4 T cells and cytotoxic intraepithelial CD8+ T cells are thought to be central in disease pathology^1,6-8^, however, CeD is a complex immune-mediated disorder and to date the findings are mostly based on analysis of heterogeneous cell populations and on animal models. Here, we comprehensively explore the cellular heterogeneity of CD45+ immune cells in human small intestine using index-sorting single-cell RNA-sequencing^9,10^. We find that myeloid and mast cell transcriptomes are reshaped in CeD. We observe extensive changes in the proportion and transcriptomes of CD4+ and CD8+ T cells and define a CD3zeta expressing NK-T-like cell population present in the control LP and epithelial layers that is absent and replaced in CeD. Our findings show that the immune landscape is dramatically changed in active CeD which provide new insights and considerably extend the current knowledge of CeD immunopathology.

---

To assess the changes in the immune cell landscape of the small intestine in CeD, we isolated CD45^+^ immune cells from LP and the epithelial layer of duodenal biopsies obtained from newly diagnosed CeD patients and from individuals with normal histology (controls) (Fig. 1a and Supplementary Table 1). In line with previous studies^11,12^, analysis of the markers used for index sorting shows that CeD harbors increased numbers of CD45+ cells in LP and a higher fraction of CD3+ intraepithelial lymphocytes (IELs) as compared to controls (Extended Data Fig. 1a, b). A modified SORT-seq^10^ single-cell RNA-seq protocol and application of quality control cut offs yielded ∼4,000 single cells with high quality RNA profiles (Fig. 1a and Extended Data Fig. 1c-e). t-Distributed Stochastic Neighbor Embedding (t-SNE) analysis revealed five major cell compartments that, based on typical lineage markers (assessed by protein and/or mRNA), were annotated as T-cells (CD3), plasma cells (PCs) (CD19 and *SDC1* (CD138)), myeloid cells (CD14, CD11c and HLA-DR), mast cells (MC) (*KIT*) and B cells (*MS4A1* (CD20)) (Fig. 1b-d). Cell size and granularity are in line with the classical phenotype of the cells (Fig. 1e). The tSNE clusters were formed by contribution from all donors with no considerable donor batch effect (Extended Data Fig. 1f).

**Fig. 1.**
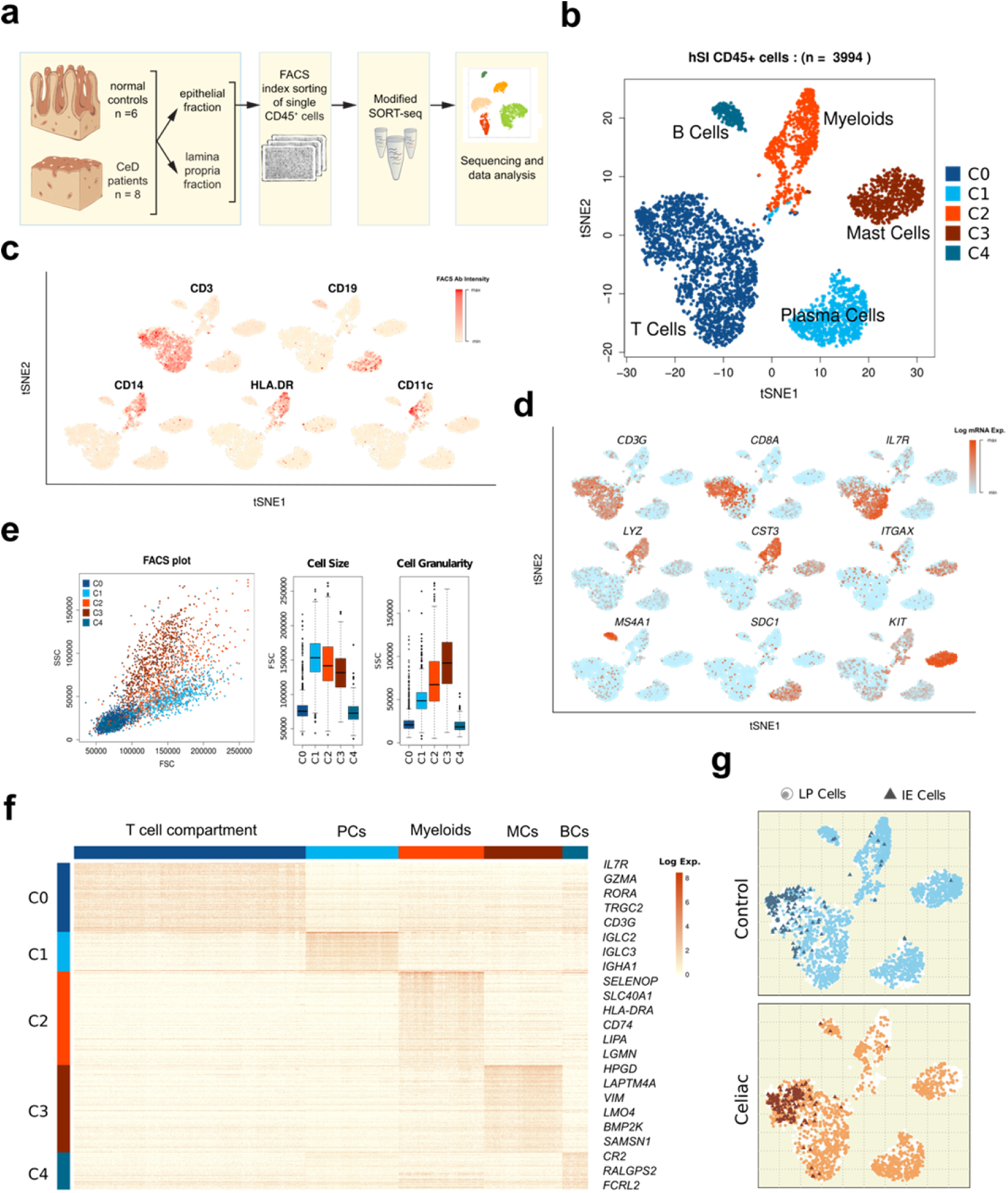
Single-cell CD45+ immune cells in human small intestine (hSI) of CeD and control samples. **a**, Schematic study design **b**, tSNE cluster plot of hSI CD45+ cells based on single cell mRNA expressions. **c**, Flowcytometric antibody intensity plots of the canonical markers in the CD45+ single cells in CeD and normal controls. **d**, Single-cell RNA-seq mRNA expression of selected genes in the tSNE-clusters of the CD45+ cells. **e**, Flowcytometric analysis scatter and box plots of CD45+ cells using FSC and SSC read outs. **f**, Heatmap of differentially expressed genes of each cluster (cell type) showing the top ranked genes. **g**, Projection of the Celiac and control derived cells on the tSNE clusters and the IELs in Celiac and control samples.

We applied the Wilcoxon rank sum test (‘Seurat’ pipeline^13^) to identify cell type-specific transcripts distinguishing each cell population (Fig 1f and Supplementary Table 2). We identified 181 genes differentially expressed in the T-cell compartment with *IL7R, GZMA, RORA* amongst the top genes. For PCs, 104 genes largely consisting of immunoglobulin related genes such as *IGLC2, IGLC3*, and *IGHA1.* We found 245 marker genes distinguishing myeloid cells with *SELENOP, SLC40A1, HLA*-*DRA* and *CD74* among the top genes. MC analysis yielded 229 genes with the top ranked genes being *HPGD, LAPTM4A, VIM* and *LMO4*. We could identify in total 98 genes as B cell markers with *CR2, RALGPS2* and *FCRL2* as top ranked genes (Fig. 1f). CeD and control cells populate the same major cell compartments but a different pattern within the MC, myeloid and T cell compartments was observed. Virtually all CeD and control IELs clustered with the T-cell compartment (Fig. 1g).

tSNE analysis of the MC compartment revealed 4 clusters (C0-C3). C1 is common to controls and CeD, whereas C2 and C3 consist mostly of control and CeD cells, respectively (Figs. 2a, b). MCs from each donor contributed to these three clusters except for C0, which is donor-specific and was excluded from the subsequent analysis (Extended Data Fig. 2a). Cluster C3 cells, enriched in CeD, were smaller and less granular (Fig. 2c). The three clusters differ in their expression levels of *C20orf194* and *FKBP11* that are high in the common cluster C1, whereas *REG1A* and *HLA-DRB1* are high in the control cell cluster C3 and low in the CeD cluster C3 (Fig. 2d and Supplementary Table 3).

**Fig. 2.**
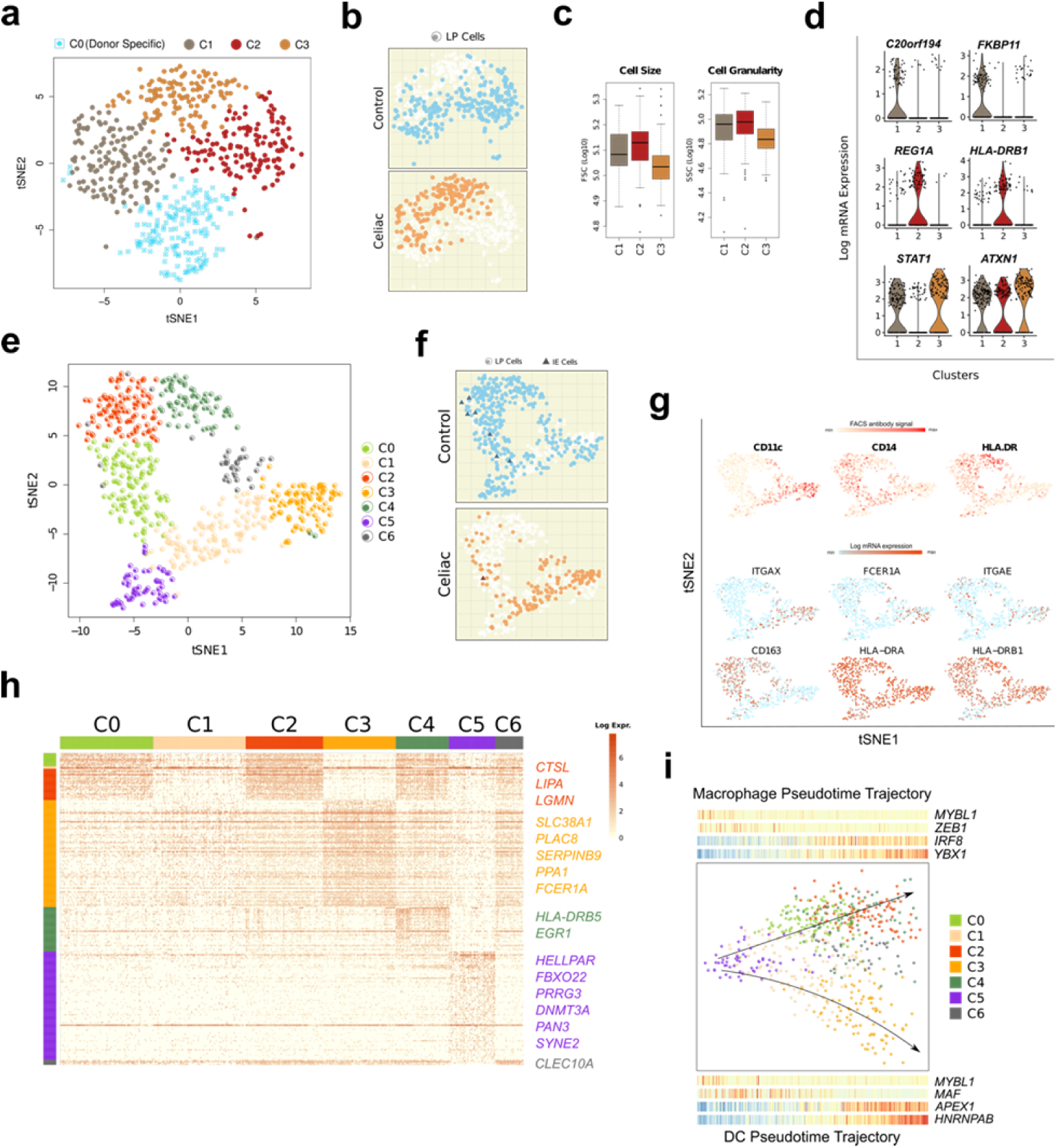
Single-cell landscape of Mast and myeloid cells in hSI of CeD and control samples. **a**, tSNE cluster plot of CD45+ mast cells in CeD and control samples. **b**, Overlay of Celiac and control cells on the mast cells tSNE clusters. **c**, Flowcytometric analysis box plots of the mast cells using FSC and SSC. **d**, Violin plots of selected mast cell cluster marker genes. **e**, tSNE clustering plot of mRNA expressions of CD45+ myeloid cells in Celiac and control samples. **f**, Projection of Celiac and control cells from Intraepithelium and Lamina Propria cells on the myeloid cell tSNE clusters. **g**, Flowcytometric antibody intensity of CD11c, CD14 and HLA-DR proteins in the myeloid cells on the tSNE clusters (top) and mRNA expression patterns of selected genes across the myeloid tSNE clusters (bottom). **h**, Heatmap of differentially expressed genes of myeloid clusters depicting the selected top ranked genes. **i**, Pseudotime analysis plots of the myeloid cells with corresponding ‘Macrophage’ and ‘DC’ trajectory scatter plots depicted underneath the top expressed TFs in bins 1 to 5 of the trajectory path (side legends correspond to the myeloid tSNE-clusters involved in each trajectory path).

tSNE analysis of the myeloid compartment revealed seven clusters (C0-C6) (Fig. 2e). Clusters C0, C2, and C4 express higher levels of CD163 and lower levels of CD11c (Fig 2e, g). C2 and C4 differentially express typical macrophage genes^14^ (Fig. 2h and Supplementary Table 3) reminiscent of CD11c-mature macrophages (Mf) which are noticeably underrepresented in CeD^15^ (Fig. 2f and Extended Data Fig. 2b, c). Clusters C1 and C3 express higher levels of ITGAX (CD11c) and FCER1A reminiscent of immature Mf and of dendritic cells (DC)^16^ (Fig. 2e, g). CD11c-high cells were ordered from C1 to C3 and C6 cells with C3 cells differentially expressing *SLC38A1, PLAC8, SERPINB9, PPA1* and *FCER1A* and C6 cells expressing *CLEC10A*, a marker associated with cDC2^17^ (Fig. 2h). Pseudotime analysis revealed two trajectories originating from cluster C5 and diverging into a Mf and a DC path that correspond to CD11c-low and CD11c-high cells, respectively (Fig. 2i). C6 cells expressing *CLEC10A* are located between the two trajectories suggesting that this cluster may originate from both the Mf and the DC pathway^16^. C5 cells differentially express *HELLPAR, FBXO22* and *PRRG3* (Fig. 2h). Note that the fluorescent antibody panel used in the index sorting cannot distinguish C5 cells from the other myeloid cells (Extended Data Fig. 2d, e). Using linear modeling and Bayesian statistics, we identified *MYBL1* as a highly expressed transcription factor (TF) in the shared origin of Mf and DC (C5) with *ZEB1, IRF8* and *YBX1* as the potential driver TFs of the MF path and *MAF* of the DC path (Fig. 2i and Supplementary Table 4).

Focusing on the humeral branch of the adaptive immune system, tSNE analysis identified a small cluster of B cells (C2) and two prominent clusters (C0 and C1) of PCs (Fig. 3a). FACS analysis showed that CD19+ PCs are significantly increased in CeD (Extended Data Fig. 3a). Based on transcriptome analysis, we noticed that B cells and PCs are rather equally distributed across the clusters in CeD and control samples (Fig. 3b and Extended Data Fig. 3b, c). Beside subtle differences in immunoglobulin genes, no major difference between the two PC clusters could be detected in mRNA, CD19 protein expression and cell size and granularity (Extended Data Fig. 3d-f and Supplementary Table 5). Pseudotime analysis revealed B cells at the beginning of the path and the two PCs clusters largely intermingled along the trajectory (Fig. 3c). Using our linear modeling approach, we identified 1597 and 1210 genes that are gradually up- or down-regulated (Supplementary Table 6). The TFs *ETS1, BCL11A, REL*, and *IRF8* are found to be highly expressed in B cells and their expression is downregulated in PCs. *MEIS2* and *RUNX1* appear to be transiently upregulated (bin 3) along the pseudotime path whereas *XBP1, CREB3L2, SUB1*and *PRDM1* are among the highest expressed TFs in fully differentiated PCs (Fig. 3d and Supplementary Table 6).

**Fig. 3.**
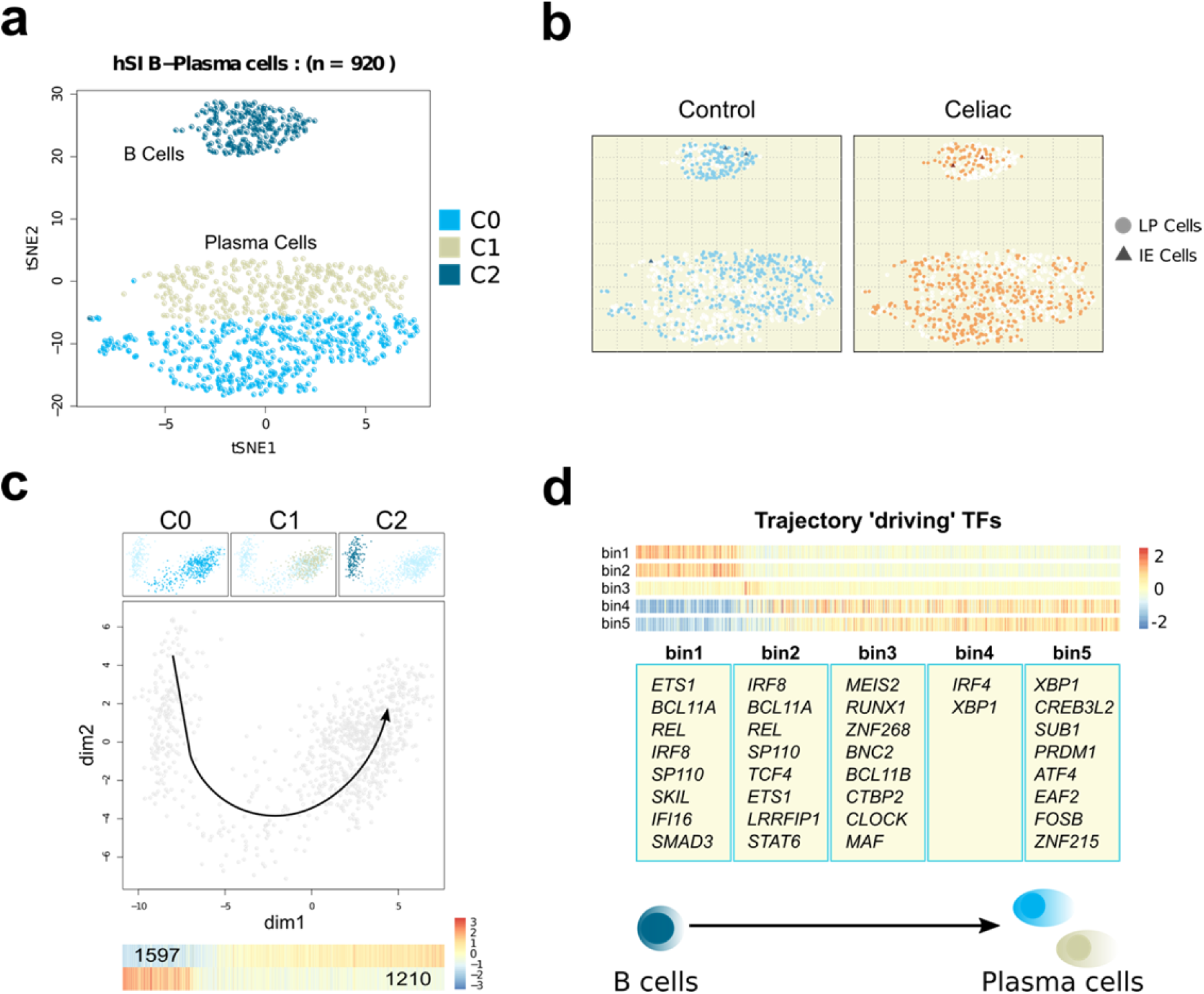
Single-cell landscape of B and plasma cells in hSI in CeD and control samples. **a**, tSNE analysis clustering plot of the B and Plasma cells in Celiac and control using single-cell RNA-seq expression profiles. **b**, Overlay of Celiac and control derived cells from Intraepithelium and LP on on the tSNE-clusters. **c**, Pseudotime analysis plot of B and Plasma cells with projection of the tSNE clusters (top scatter plots) and heatmap of up- and down-regulated genes along the B to Plasma cells trajectory path. **d**, Heatmaps and the top ranked potential ‘driving’ TFs obtained by the linear modeling that are differentially expressed in each bin of the pseudotime trajectory along the B to Plasma cell paths.

tSNE analysis of the T-cell compartment (Fig. 1b) yielded three clusters of *CD4*^+^ T cells (C0, C2 and C4) and four clusters of *CD8*^+^ T cells (C1, C3, C6 and C7) (Fig. 4a and Extended Data Fig. 4a). Three clusters (C5, C8 and C9), that co-cluster with the T-cell compartment when analyzing all CD45^+^ cells (Fig 1), turn out to be distinct and are CD4^-^ /CD8^-^at mRNA levels and express no or very low levels of CD3 protein (Fig. 4a-c and Extended Data Fig. 4c). Expression of the canonical *GNLY* ^*18*^ amongst 29 other known and novel genes, identified cluster C8 as natural killer (NK) cells (Fig. 4e and Supplementary Table 7). The NK cells are mainly present in the LP and more abundant in CeD than controls (Fig. 4 d, g). Cells of cluster C9 were less present in CeD and identified as innate lymphoid cells class 3 (ILC3) based on their expression of *PCDH9, KIT, ALDOC, AHR* and *RUNX2* genes as previously reported^19^ (Figs. 4a, d, e and Supplementary Table 7).

**Fig. 4.**
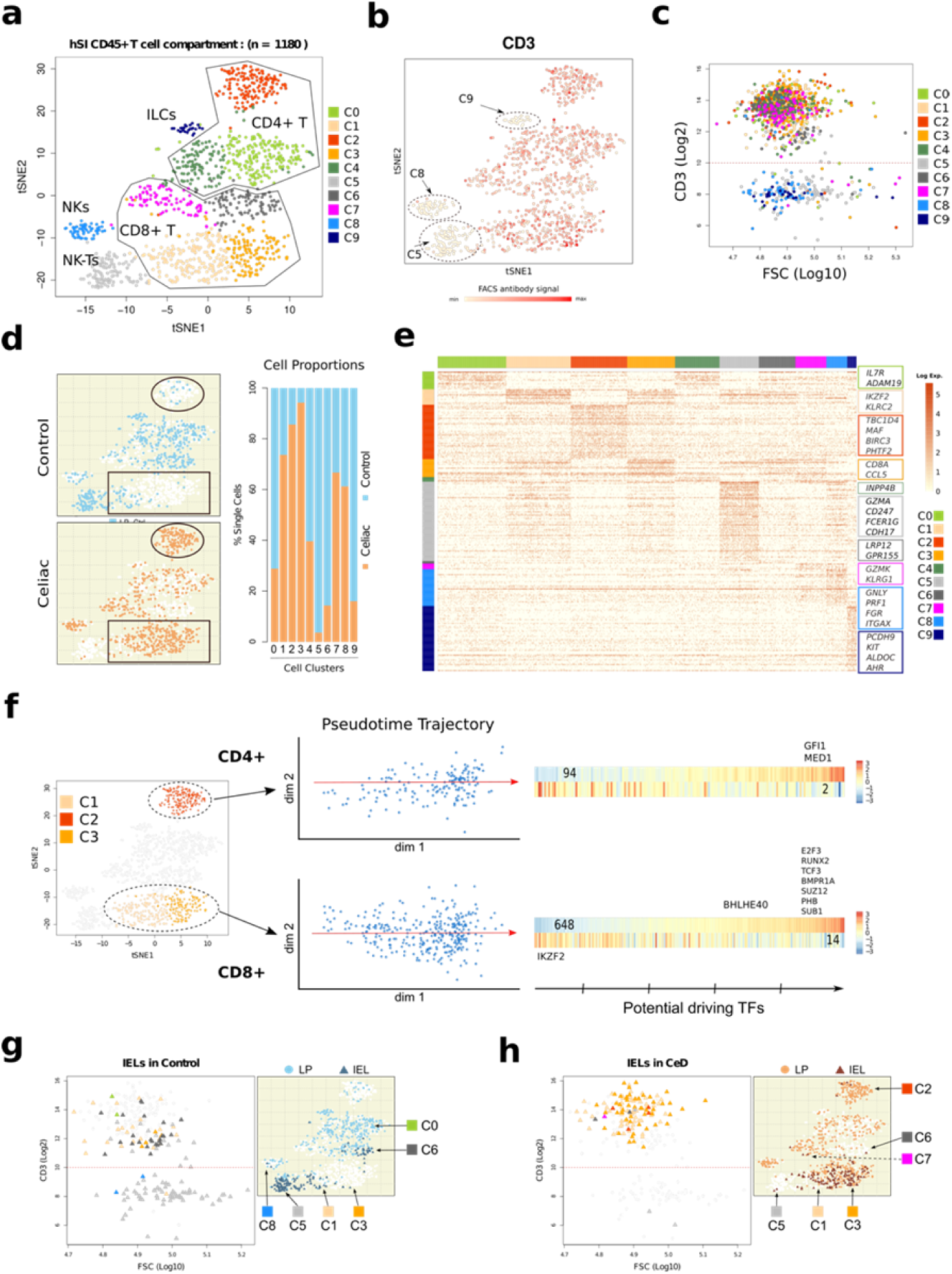
Single-cell landscape of T-cell compartment in hSI of CeD and control samples. **a**, tSNE clustering plot of the cells in T-cell compartment (as in Figure 2a) using single-cell RNA-seq transcriptome profiles depicting *CD4*+, *CD8*+, ILCs, NK and NK-T cell clusters. **b**, Flowcytometric analysis scatter plot of CD3 protein expression in the T-cell compartment across related tSNE clusters. **c**, Scatter plot of CD3 expression and FSC signals showing CD3high and Cd3low cells. **d**, Projection of Celiac and control derived T-cell compartment cells showing the CeD CD8+ enriched (rectangles) and CeD CD4+ enriched (ovals) clusters together with the related proportional bar plots on the tSNE-resulted clusters. **e**, Heatmap of differentially expressed genes among tSNE-clusters showing the top ranked genes for each cluster. **f**, Pseudotime analysis scatter plots of the CD4+ (top panel) and CD8+ (bottom panel) cells depicting the trajectory paths and the related heatmaps of up- and down-regulated genes along each path with the top potential ‘driving’ TFs present in each bin (bottom axis; five bin segments) along the trajectory. **g, h**, Scatter plots of FACS CD3 expression (left) and tSNE cluster distribution (right) of IELs in control samples and in CeD, respectively.

Strikingly, the CD4+ T cells in CeD are different from controls. While control cells dominate cluster C0 expressing among other genes *IL7R, ADAM19, TNFSF13*, more than 85% of cells in C2 are from CeD (Fig. 4d). C2 cells express genes including *TBC1D4, MAF, BIRC3, PHTF2, TFRC, TIGIT, IL32, FOXO1, GBP5, STAT1*, and *IL6R*; several of these are known to be expressed in both activated and regulatory T cells ^*20-22*^ (Fig. 4e and Supplementary Table 7). Gene ontology analysis showed that these cells are significantly enriched for signaling pathways including ‘Cytokine Signaling in Immune system’, ‘NOD-like receptor signaling pathway’ and ‘Signaling by Interleukins’ (Supplementary Table 7). Pseudotime analysis on C2 cells revealed 94 genes that are upregulated along the trajectory with *GFI1* and *MED1* as the highest expressed TFs in the end of the path (Fig. 4f). *GFI1* and *MED1* have been shown to regulate T cell activation and NKT cell development, respectively^23,24^. C4 cells are equally present in CeD and controls and express genes including *INPP4B, ANK3, FOXP1* and *AHNAK* (Figs. 4d, e).

CD8^+^ T cells in CeD clusters are also very differently from controls. Clusters C1 and C3 contain 75% and 95% CeD cells, respectively (Fig. 4d). Importantly, the clusters contain the vast majority of both LP and intraepithelial CD8+ T cells (Fig. 4h). C1 cells differentially express *KLRC2* and *KLRD1* both NK markers known to be involved in the cytotoxic activity of CD8 IELs in CeD^25,26^. They also express *TRGC2* suggesting that gamma delta T cells are within this cluster. C3 cells express *ITGAE* and *CD101*, marker genes for resident memory T cells^27^, and *CCL5, HLA-DP* and *ENTPD1* (CD39), all expressed on activated CD8+ T cells (Fig. 4e and Supplementary Table 7). Interestingly, CD39 was recently shown to be a marker for tumor-specific CD8 T cells^28^. Control CD8^+^ T cells mainly contribute to cluster C6 and highly express *LRP12* and *GPR155*. Cluster C7 contains slightly more CeD cells that differentially express *GZMK* and *KLRG1* suggesting they are recently recruited memory CD8^+^ T cells^29^ (Figs. 4a, d, e and Extended Data Fig. 4c and Supplementary Table 7). Pseudotime analysis of clusters C1 and C3 revealed a developmental trajectory in which it appears that C1 cells progress towards C3 cells. Applying our linear modeling, we found that *IKZF2* is the potential driving TF expressed in C1 cells and *E2F3, RUNX2*, and *TCF3* among others in C3 (Fig. 4f and Supplementary Table 8).

Our analysis reveals that IELs in control individuals consist of two distinct populations: CD3low and CD3high cells. The CD3high IELs are mostly CD8^+^ T cells (Figs. 4g). The CD3low IELs are bigger in size and more granular as compared to the other cells in the T-cell compartment (Extended Data Fig. 4b) and are mainly CD4low/CD8low/ CD247 high (CD3 zeta) cells (cluster C5, Figs. 4a, b, g and Extended Data Fig. 4c). We classify cluster C5 cells as ‘NK-T’ as they express classical NK as well as T cell genes such as *TRDC, GZMA, FCER1G, CDH17, CD69, TYROBP* and *ID2* (Fig. 4e and Supplementary Table 7). This transcriptome profile is very similar to that described for natural IELs in mice^30^. The CD3low cells in C5 are not exclusively intraepithelial as ∼25% of the cells are located in the LP (Extended Data Fig. 4d). Moreover, C5 cells are remarkably underrepresented or absent in CeD (Figs. 4d, h and Extended Data Fig. 4d). Together, we find that the majority of CD8+ T cells in both LP and epithelium express an activated phenotype in CeD compared to controls, suggesting that CD8+ T cells in both compartments are actively involved in CeD pathology.

In conclusion, we find a dramatic change of the immune cell landscape in the active CeD lesion. The number of CD3+ T cells is increased and the majority of CD4+ and CD8+ T cells are transcriptionally very different from their control counterparts. At the same time mature Mf with regulatory properties are greatly reduced and NK-T IELs are virtually absent in CeD. Our atlas thus provides an important framework to increase our knowledge about this complex immune-mediated disease.

## Acknowledgements

HGS and NA are supported by the Central European University European Research Council Advanced Grant SysStemCell (339431).

## Author Contributions

NA, AB, ESB, FLJ and HGS designed the study. NA, EJ-M and PBB set up the experimental work flow. NA performed the single cell RNAseq experiments, analyzed the single cell transcriptome data and integrated the flowcytometry data. AB and ESB performed the single cell isolation, antibody indexing and FACS sorting and analyzed the flowcytometry data. JJ responsible for patient material. All authors contributed to the writing of the manuscript. HGS is the corresponding author who supervised the study and contributed in data interpretations.

## Additional information

**Extended data** is available for this study at <u>http://</u>

**Supplementary information** is available for this study at <u>http://</u>

## Extended data

### Methods

#### Tissue samples

Duodenal biopsies were obtained during routine endoscopy at Akershus University Hospital from patients referred due to suspicion of celiac disease. Diagnosis of celiac disease followed standard procedure (Ludvigsson et al., 2014). 8 (all females, median age 29.7) patients with confirmed untreated celiac disease (Marsh score 3A-3C) and 6 (5 females, median age 35.8) confirmed non-celiac controls have been enrolled in the study. See Table 1 for the list of patients and their clinical information. Biopsies have been placed immediately in ice-cold buffered saline, transported on ice, and processed within 4h. The study was approved by the Norwegian Regional Committee for Medical Research ethics (REK 2012/341), and all patients gave written informed consent.

#### Preparation of single cells suspension

To separate the epithelium and intraepithelial lymphocytes biopsies were shaken twice in 6.5 ml of PBS with 2mM EDTA (Sigma-Aldrich), 1% FCS (Sigma-Aldrich) and 1µM flavopirydol (Sigma-Aldrich) for 10 min at 37°C. Supernatants containing the epithelial fractions were combined, washed, passed through 100 µm cell strainers (Miltenyi Biotec), washed again and kept on ice until staining. Epithelium-free mucosa was minced and incubated with stirring in 2.5 ml of RPMI1640 (Lonza) containing 10% FCS, 1% Pen/Strep (Lonza), 1µM flavopirydol, 0.25 mg/ml Liberase TL (Roche) and 20 U/ml DNase I (Sigma) for 40 min at 37°C. Halfway through the incubation the samples were triturated using 1 ml pipette to facilitate complete digestion. Digested cell suspension was passed through 100 µm cell strainer and washed twice.

#### FACS sorting

Cells were stained with a combination of fluorescent antibodies in FACS buffer for 20 min on ice, washed twice, and filtered through 100 µm nylon filter mesh before sorting. Both lamina propria and epithelial fractions were stained with: FcR Blocking Reagent (Miltenyi Biotec), CD45-APC-H7 (2D1, BD Biosciences), CD3-APC (OKT3, Biolegend), CD19-BV421 (HIB19, Biolegend), HLA-DR-PerCP-Cy5.5 (L234, Biolegend), CD14-PE-Cy7 (HCD14, Biolegend), CD11c-PE (S-HCL-3, BD Biosciences). EpCAM-FITC (Ber-EP4, Dako) was added to the epithelial fraction to label epithelial cells. In some samples CD27-BV605 (O323, Biolegend) and CD103-BV605 (Ber-ACT8, Biolegend) were added to the lamina propria and epithelium fraction, respectively. Dead cells were stained and excluded using To-Pro-1 (Molecular Probes). Doublets were identified and excluded based on scatter parameters in forward scatter area versus height (FSC-A/FSC-H) and side scatter area versus width (SSC-A/SSC-W) plots. Cells of interest (single live CD45+, or single live CD45+(CD27-)CD3-CD19- to enrich for myeloid cells) were index sorted into Bio-Rad Hard Shell 384 well microplates (Bio-Rad) containing well specific primers (100 nl, 0.75pmol/ul) and 5 ul mineral oil (Sigma-Aldrich) on top using FACS Aria IIu or FACS Aria III with FACS Diva software version 8 (BD Biosciences). After sorting plates were sealed, labelled, span down for 10 min at 2000g, snap frozen on dry ice and kept at −80°C until used. In addition, a minimum of 10^5^ total events from each sample were recorded and exported with index sort files for analysis in FlowJo software.

#### scRNA-seq modified SORT-seq

In order to obtain the transcriptome of immune cells of small intestine in Celiac and normal controls, we used the SORTseq protocol (Muraro *et al*. 2016) which itself is a derivates of original CEL-Seq2 protocol (Hashimshony, T. et al 2016) and applied some modifications to the original protocols. In brief, single immune cells were first indexed with canonical markers of interest using the previously described antibodies and then sorted in 384-well PCR plates (BioRad). The plates were already primed with unique primers per each well in which each primer has a unique cell barcode with UMI incorporated to tag each RNA molecule in the cell. Sorted plates then were kept frozen at −80 °C before proceeding to cDNA synthesis. The frozen plates were shortly centrifuged at 1200 RPM for 1 minute at 4 °C and incubated in the PCR machine at 60 °C for 2 minutes to break the cells and release the RNA.

We used micro-dispenser machine, Nanodrop II (BioNex) to dispend all the reagents in the next steps of library preparation. After cell breakage, 100 nl lysis buffer contains RNase inhibitor (RNasin PLUS promega), Triton X100, ERCC spike in control RNAs 1:50,000 diluted (Ambion) were added to each well, short spinned at 1200 RPM for 1 minute and incubated in the PCR machine at 65 °C for 2 minutes and then immediately put on ice for 5 minutes.

Then first 150 nl cDNA reaction mix contains first strand cDNA synthesis buffer (Invitrogen), SuperscriptII reverse transcriptase (Invitrogen), 0.1M DTT and RNasein PLUS (promega) was added to each well of the plate and the plates were spinned at 1200 RPM at 4 °C for 1 minute and placed in PCR machine at 42 °C for 1 hour followed by 10 minutes enzyme deactivation at 70 °C and immediately placed on ice.

Then, the 500 nl second strand DNA reaction mix contains second strand cDNA synthesis buffer (Invitrogen), 10 mM dNTP, E.Coli DNA ligase enzyme (NEB), E.coli DNA polymerase enzyme (NEB) and RNaseH enzyme (com) was added to each well of the plate and then plates were spinned for 1 minute at 1200 RPM and placed in the PCR machines at 16 °C for 2 hours followed by enzyme deactivation at 70 °C for 10 minutes and then placed on ice.

To remove the excess remainder of the primers in the reaction mix in each well, we used exonuclease-I enzyme (NEB) and incubated the plates at 37 °C for 20 minutes followed by enzyme deactivation at 85 °C for 15 minutes and immediately placed the plates on ice. To reduce the DNA loss during the subsequent steps of library preparation, we linearly amplified the cDNA content of each well using unidirectional linear-PCR using Pfu DNA polymerase enzyme (Promega), forward primer against the common sequence of the T7-P5 part of the original oligo-dT primers and ran the linear-PCR for 15 cycles with 2 minutes preheating at 95°C, then sequential cycles of 30 seconds at 94°C, 30 seconds at 50°C and 3 minutes at 72°C and ended by incubation at 72°C for 5 minutes and then placed plates on ice. After linear amplification of the DNA content in each well, the reaction solution of all wells per each plate were pooled together (∼ 600 ul) and the DNA was cleaned up using AMPure XP beads by mixing 100 ul XP beads with 600 ul clean up buffer contains PEG and tween and adding it to the DNA pool. The DNA-bond beads then washed two times with 80% ethanol and the DNA was eluted in 22 ul EB buffer.

Then the amplified single strand DNA fragments of each pool was second stranded using second strand DNA synthesis buffer (NEB), first strand cDNA synthesis buffer(NEB), 0.1M DTT, random hexamer primer mix (Thermo), dNTP mix, E.coli DNA polymerase (NEB) and E.coli DNA ligase (NEB) in 50 ul reaction volume and incubated for 2 hours at 16°C. Then the DNA was cleaned up using 2x XP-bead:DNA ratio and eluted in 7 ul water.

Then the DNA was amplified and transformed to RNA using in vitro transcription overnight according to manufacturer’s protocol (Ambion Mega-Script IVT kit) at 37°C. The amplified RNA (aRNA) then was treated with ExoI-rSAP enzymes for 20 minutes at 37°C and fragmented using Manganese based fragmentation buffer for 1.5 minutes at 94°C followed by incubation with 0.5M EDTA buffer to stop the fragmentation process. The fragmented aRNA was subsequently cleaned up using 2x XP-bead:aRNA ratio and eluted in 12 ul water.

We used 5 ul of the aRNA and proceeded to next step of library preparation. The cleaned aRNA was converted to cDNA using 1 ul of random octamer primers that contain part of T7 sequence of Illumine library oligos and the reaction mix contains first strand cDNA buffer (NEB), 0.1M DTT, 10mM dNTPs, RNasein PLUS (Promega) and SuperscriptII reverse transcriptase enzyme (Invitrogen) was incubated at 25°C for 10 minutes followed by incubation at 42°C for 1 hour and 70°C for 10 minutes and then placed on ice. Then the final library was amplified and enriched by 7 cycles of PCR using KAPA hyper prep kit and adding forward and reverse primers against Illumine T5 and T7 sequences in which the reverse primer contained unique sequence and added specifically per each sample/library. The library then was cleaned with 1:1 XP-bead:DNA ratio and eluted in 20 ul elution buffer. The quantity of the library was measured using KAPA library quantification kit (Roche) and the quality was checked by using the bioanalyzer machine (Agilent).

#### scRNA-seq data analysis

Each library obtained from one plate was sequenced for on average 40 million reads using NextSeq 500 sequencer. The raw FASTQ files (Read2 contains 8nt cell barcode + 8nt UMI; Read1 contains mRNA sequence) were used as input for the CEL-Seq2 pipeline (Hashimshony, T. *et al.* 2016) however, incorporating STAR aligner (https://github.com/alexdobin/STAR) to map the sequenced reads against the human genome HG38 using the default parameters. The uniquely mapped reads then were demultiplexed per each single cell per library using the known 384 cell barcodes. The demultiplexed, uniquely mapped reads per each single cell then were counted against the reference genome and PCR deduplicated using HTSeq-count part of the CEL-Seq2 pipeline and the table of read counts for all single cells (∼12,000 single cells) was built using R environment.

To remove the low-quality cells from the dataset, we applied several criteria’s. Cells with MAD of ERCC and mitochondrial counts and library size greater than 3 were removed. Next, cells with total detected genes less than 400 and more than 4000 were also excluded from the dataset. We included no-cell and 12x-cell wells during the sorting of the cells in the plates and used these wells as background and multicell controls for the analysis respectively. Cells with Pearson correlation greater than r > 0.8 as compared to the multicell controls, were excluded from the dataset as doublets. We also removed the drop-out genes from the dataset and also the genes which had expression ratio of background to signal greater than 1. The dataset of filtered high quality cells (∼4000) then was normalized using DESeq package incorporating the ERCC spike-ins and used for further analysis.

Subsequently, we used Seurat v.2 pipeline (Butler *et al.* 2018) to identify the possible clusters of cells and the main cell populations using the Wilcoxon Ranked-sum test with the default parameters. Then using Seurat, we identified marker genes per each cluster of cells and used the obtained marker genes to further do the heatmap analysis and the gene ontology analysis accordingly in R environment using pheatmap and gprofiler packages, respectively. The tSNE analysis was subsequently performed using the related integrated function within the Seurat package with the default parameters.

To assess any possible trajectory within the cells of interest, we did pseudo-time analysis on the selected cells using Slingshot algorithm incorporated in the dynverse pipeline (https://github.com/dynverse) and to check for any up-regulation and down-regulation patterns of genes within the trajectory path, we applied a linear modeling analysis incorporating a Bayesian statistics using Limma package (Ritchie ME *et al.* 2015) with the default parameters. To identify the possible driving transcription factors (TFs) along the trajectory path, we divided each of the trajectory paths into 5 bins using the start- and end-point cells along the pseudotime dominant dimension and applied the linear modeling per each bin of the trajectory path. Then the TFs with FDR < 0.05 were selected as the potential driving factors of each bin and per path. The simulation analysis was additionally performed in some examples to assess the possible difference between clusters of cells that would not result in a significant cluster markers using Seurat Wilcoxon ranked-sum algorithm and pipeline. In this regard, we randomly selected *n* number (*n=20%*total cells of the cluster*) of cells from each cluster of cells resulted from Seurat pipeline and constructed a mean-based mini-bulk sample of the related cluster of cells. We accordingly permuted this simulation for 1000 times and then applied the hierarchical clustering on the highly variable genes across the simulated mini-bulk samples and used the resulted genes for further gene ontology analysis.

## Data Availability

The dataset generated during and/or analyzed during the current study are available in the EGA database under data accession number EGAD00001005127 at https://ega-archive.org/studies/EGAS00001003751

---

**Extended Data Fig. 1.**
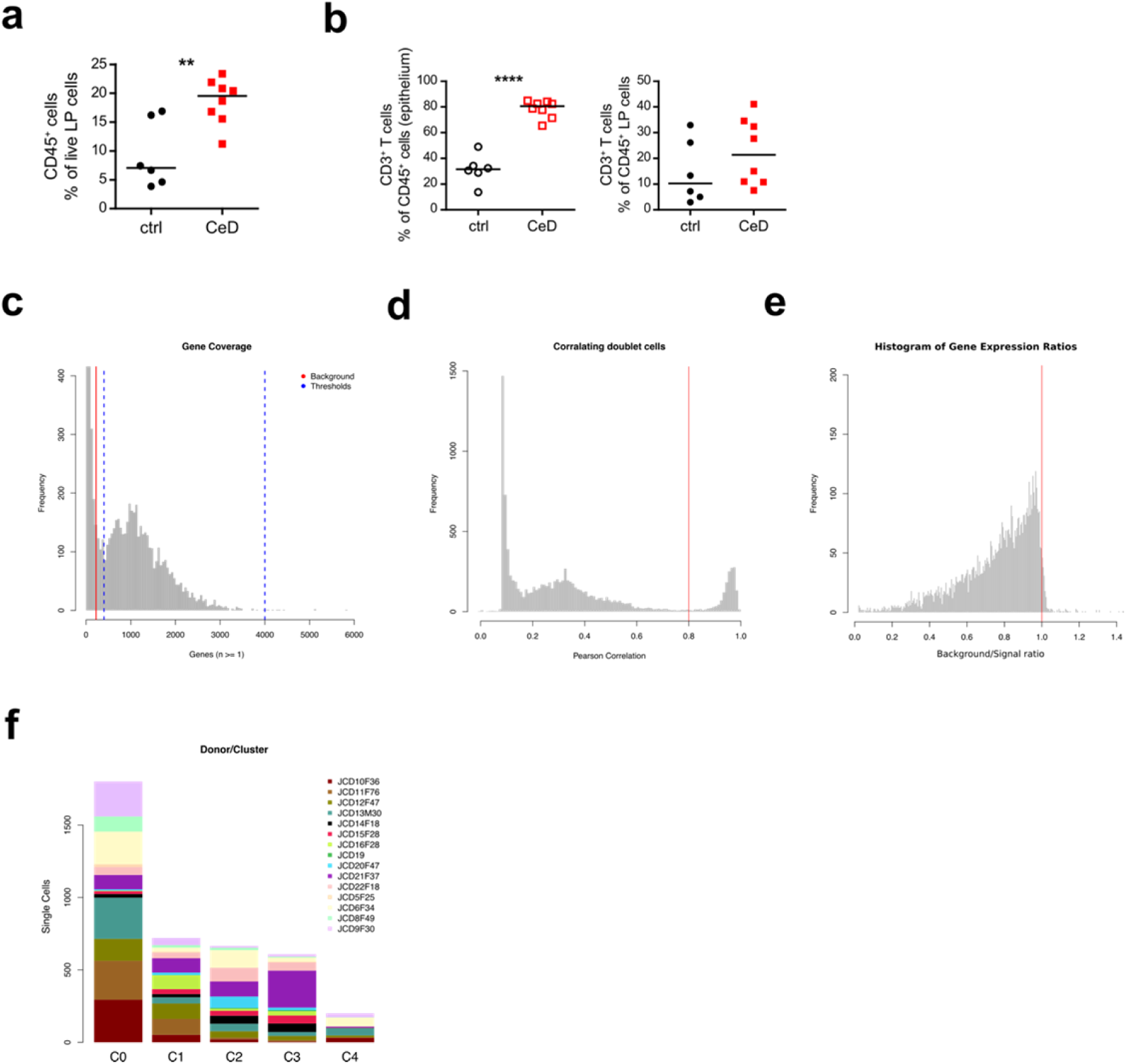
Single-cell CD45+ immune cells in human small intestine; Donor contribution and quality control. **a**, Relative representation of CD45^+^ immune cells among all live cells from control (ctrl; n = 6) and CeD (n = 8) samples. **b**, Relative representation of CD3^+^ T cells among all CD45^+^ cells from control (ctrl; n = 6) and CeD (n = 8) samples in the epithelial (left) and LP (right) fraction. **c**, Histogram plot of gene coverage per each single cell showing the average gene coverage in empty control wells as background (red line) and the applied thresholds in gene coverage (dashed blue lines). **d**, Histogram plot of Pearson correlation r scores of single cells with the control multi-cell sorted wells (vertical line showing the cut-off set to r=0,8). **e**, Histogram plot of background (empty wells) to signal (single cell sorted wells) mRNA expression ratios across all the detected genes among single cells showing the distribution of the ratios and the applied threshold; genes with background/signal ratio greater than 1.0 (red line) were removed from the data. **f**, Bar plot of donor contribution to each cluster of cells obtained from tSNE analysis.

**Extended Data Fig. 2.**
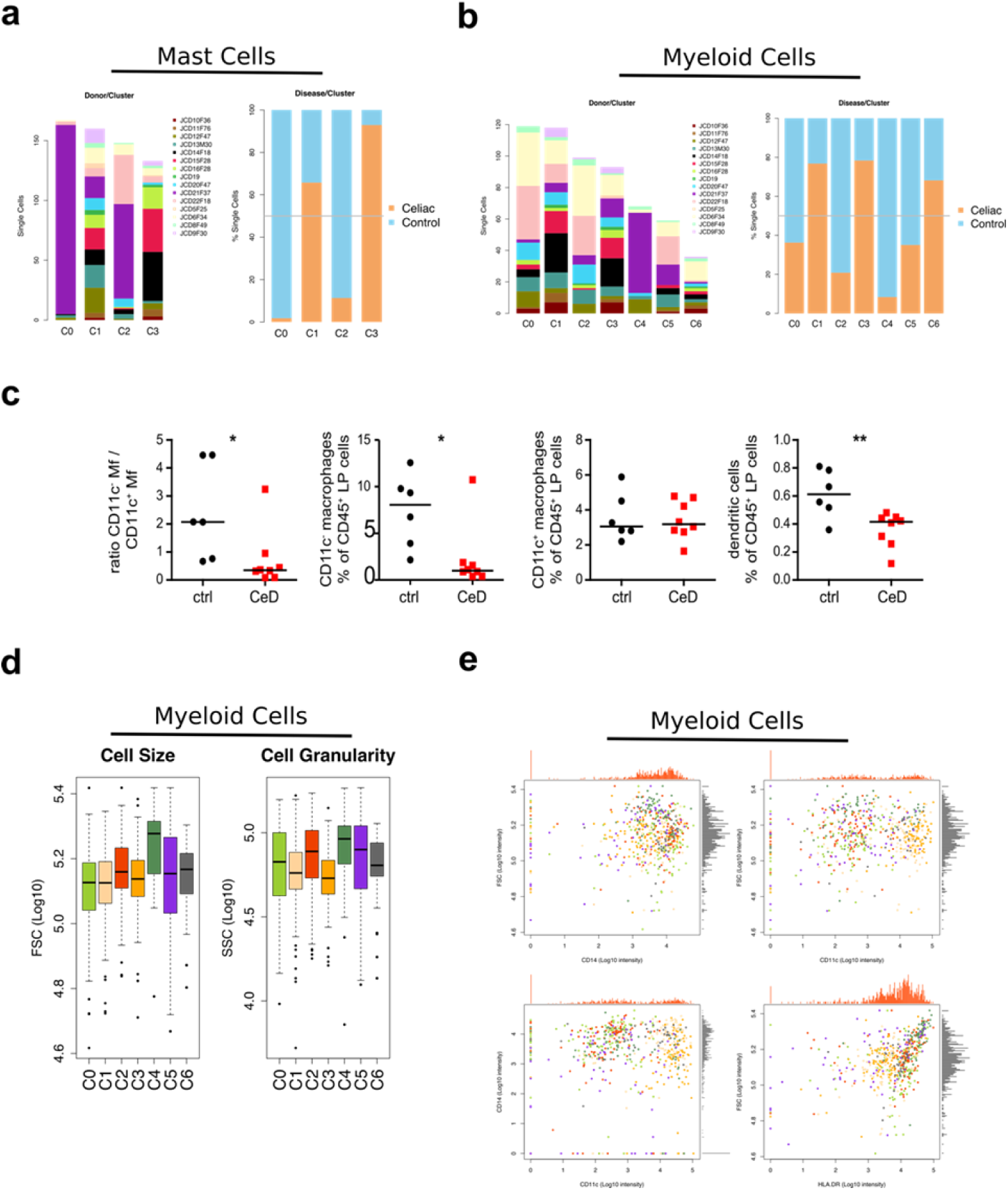
Mast and myeloid cells in hSI in CeD and control samples; FACS analysis and donor contribution. **a**, Bar plots of donor contribution in each cluster of the cells obtained from the Mast cells tSNE analysis (left) and the related Celiac vs. control contribution to each cluster (right). **b**, Bar plots of donor distribution (left) and Celiac vs. control contribution (right) per each tSNE cluster of the myeloid cells. **c**, The ratio of CD11c^-^ macrophages to CD11c^+^ macrophages (left) and relative representation of CD11c^-^ macrophages (middle left), CD11c^+^ macrophages (middle right), and dendritic cells (right) among all CD45^+^ LP cells from control (ctrl; n = 6) and CeD (n = 8) samples. **d**, Box plots of cell size and cell granularity measured via flowcytometric analysis on the myeloid cells. **e**, Flowcytometric analysis scatter plots of selected markers in the myeloid cells.

**Extended Data Fig. 3.**
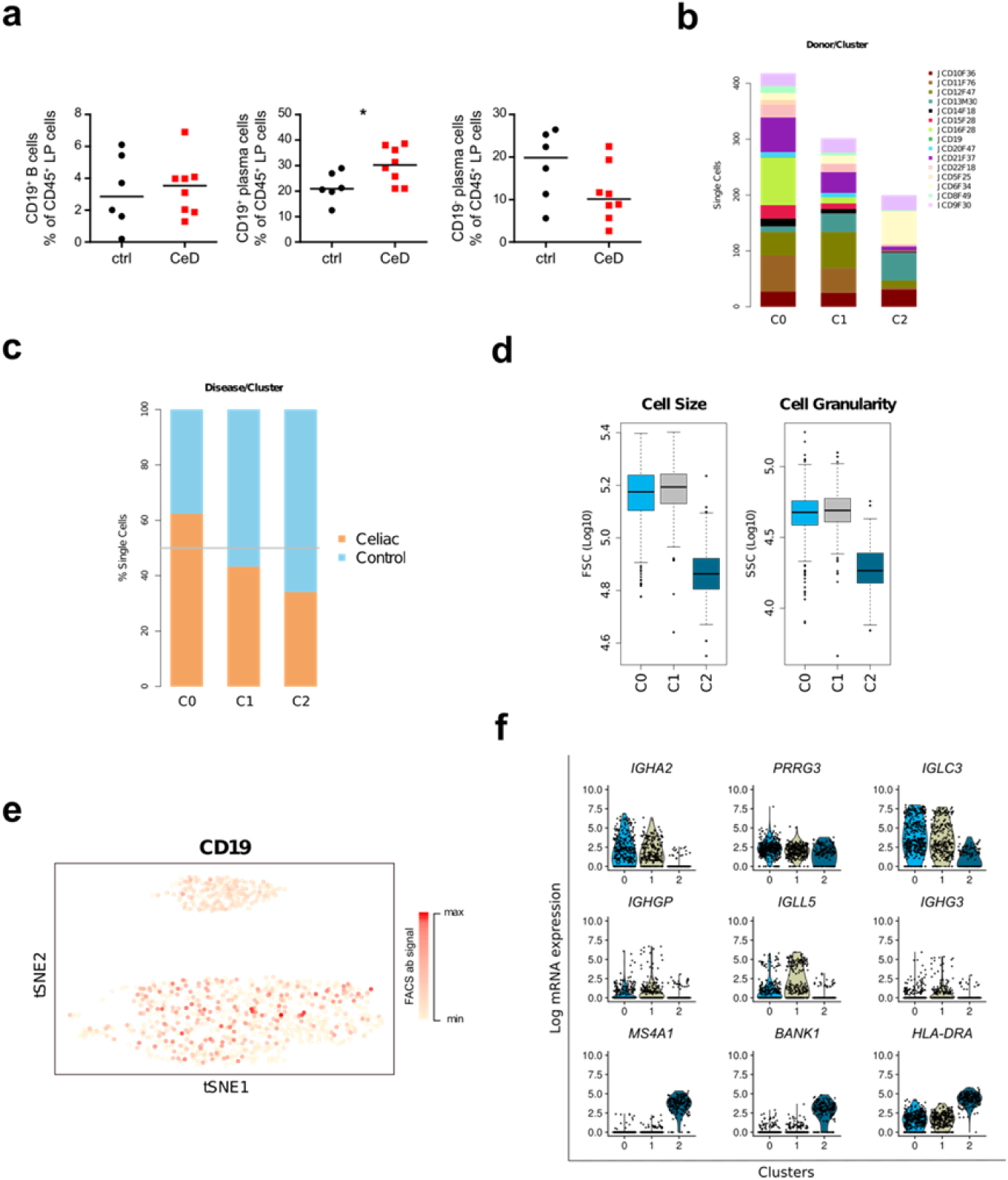
B and plasma cells in hSI in CeD and control samples. **a**, Relative representation of CD19^+^ B cells (left), CD19^+^ plasma cells (middle), and CD19^-^ plasma cells (right) among all CD45^+^ LP cells from control (ctrl; n = 6) and CeD (n = 8) samples. **b**, Donor contribution bar plot of B and Plasma cells per each tSNE derived cluster. **c**, Bar plot of Celiac vs. control contribution per each tSNE-cluster of cells. **d**, Flowcytometric analysis box plots of B and Plasma cells using FSC and SSC read outs. **e**, Scatter plot of CD19 protein expression among all B and Plasma cells on the tSNE plot using flowcytometric signal intensities. **f**, Violin plots of the differentially expressed genes between B- and Plasma-cell clusters resulted from the tSNE analysis depicting the top ranked candidate genes.

**Extended Data Fig. 4.**
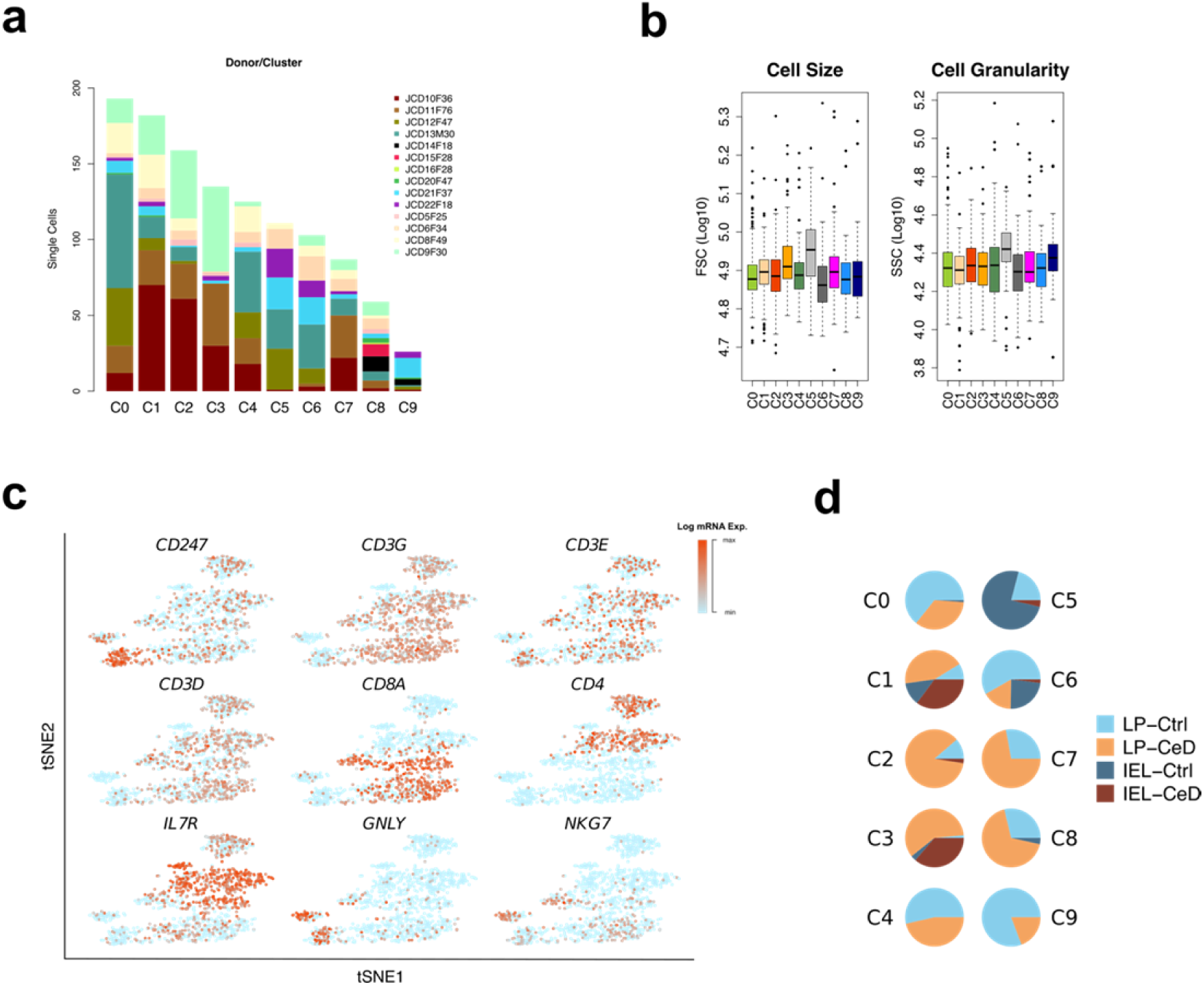
T-cell compartment in hSI in CeD and control samples. **a**, Bar plot of donor distribution per each cluster of cells obtained from tSNE analysis on the T-cell compartment. **b**, Flowcytometric analysis box plots depicting the cell size and cell granularity across all tSNE-clusters of T-cell compartment using FSC and SSC read outs **c**, Scatter plots of mRNA expression of selected genes on the tSNE clusters of T-cell compartment showing the *CD*4+ and *CD*8+ expressing clusters. **d**, Pie plots of T-cell clusters showing the relative contribution of LP and IE obtained from Celiac and control samples.

